# Clinicogenomic characterization of inflammatory breast cancer

**DOI:** 10.1101/2024.05.07.592972

**Authors:** Nolan Priedigkeit, Beth Harrison, Robert Shue, Melissa Hughes, Yvonne Li, Gregory J. Kirkner, Liam F. Spurr, Marie Claire Remolano, Sarah Strauss, Janet Files, Anne-Marie Feeney, Libby Grant, Ayesha Mohammed-Abreu, Ana Garrido-Castro, Romualdo Barroso Sousa, Brittany Bychkovsky, Faina Nakhlis, Jennifer R. Bellon, Tari A. King, Eric P. Winer, Neal Lindeman, Bruce E. Johnson, Lynette Sholl, Deborah Dillon, Beth Overmoyer, Sara M. Tolaney, Andrew Cherniack, Nancy U. Lin, Filipa Lynce

## Abstract

**Background:** Inflammatory breast cancer (IBC) is a rare and poorly characterized type of breast cancer with an aggressive clinical presentation. The biological mechanisms driving the IBC phenotype are relatively undefined—partially due to a lack of comprehensive, large-scale genomic studies and limited clinical cohorts.

**Patients and Methods:** A retrospective analysis of 2457 patients with metastatic breast cancer who underwent targeted tumor-only DNA-sequencing was performed at Dana-Farber Cancer Institute. Clinicopathologic, single nucleotide variant (SNV), copy number variant (CNV) and tumor mutational burden (TMB) comparisons were made between clinically confirmed IBC cases within a dedicated IBC center versus non-IBC cases.

**Results:** Clinicopathologic differences between IBC and non-IBC cases were consistent with prior reports—including IBC being associated with younger age at diagnosis, higher grade, and enrichment with hormone receptor (HR)-negative and HER2-positive tumors. The most frequent somatic alterations in IBC involved *TP53* (72%), *ERBB2* (32%), *PIK3CA* (24%), *CCND1* (12%), *MYC* (9%), *FGFR1* (8%) and *GATA3* (8%). A multivariate logistic regression analysis revealed a significant enrichment in *TP53* SNVs in IBC; particularly in HER2-positive and HR-positive disease which was associated with worse outcomes. Tumor mutational burden (TMB) did not differ substantially between IBC and non-IBC cases and a pathway analysis revealed an enrichment in NOTCH pathway alterations in HER2-positive disease.

**Conclusion:** Taken together, this study provides a comprehensive, clinically informed landscape of somatic alterations in a large cohort of patients with IBC. Our data support higher frequency of *TP53* mutations and a potential enrichment in NOTCH pathway activation—but overall; a lack of major genomic differences. These results both reinforce the importance of *TP53* alterations in IBC pathogenesis as well as their influence on clinical outcomes; but also suggest additional analyses beyond somatic DNA-level changes are warranted.

## BACKGROUND

Inflammatory breast cancer (IBC) is a rare and aggressive disease with unique histopathological and clinical behaviors^1, 2^. Patients with IBC have worse outcomes^3^, are enriched for the more proliferative clinical subtypes of breast cancer (triple-negative and HER2-positive disease)^4, 5^, more often present with de novo metastases, and experience shorter breast cancer specific survival^5, 6^. Despite this uniquely aggressive clinical presentation and distinct histopathologic features such as dermal lymphatic invasion^7^, the molecular drivers of the IBC phenotype remain poorly defined. To improve research outcomes and develop more biologically informed and effective therapies, it is imperative to conduct more thorough molecular analyses of this rare yet deadly disease.

The genomics of IBC has been interrogated by a few studies, yet the rarity of the diagnosis—estimated to be 1-2% of breast cancers^8^—makes large-scale analyses difficult. Nonetheless, efforts have attempted to identify pathways and genomic alterations specific to IBC with potential enrichments in TGF-B signaling^9^ and MYC amplifications^10^, as well as a recent whole-genome sequencing study of 20 patients which showed minimal genomic differences between IBC and non-IBC cases^11^. Additionally, IBC is composed of different proportions of molecular subtypes vs non-IBC, which makes it a challenge to attribute molecular differences to true IBC biology— especially with a limited number of cases per study. Lastly, even the clinical classification of IBC can be wrought with misdiagnosis, further complicating the ability to interrogate the disease accurately and comprehensively^12^.

To address these challenges and better define clinicogenomic features enriched in IBC, we compared clinicopathological and associated targeted DNA-sequencing data from a large cohort (n = 140) of advanced IBC cases to non-IBC cases. Importantly, each IBC case was identified and reviewed within a dedicated IBC Center at a single institution and harbored associated clinical and genomic data—making this the most comprehensive correlative study of IBC to date.

## METHODS

### Institutional Review Board (IRB), Cohorts, and Inclusion Criteria

We conducted this study using a prospectively maintained institutional database with both clinicopathological and genomic data from Dana-Farber Cancer Institute (DFCI) and Brigham and Women’s Hospital. The study was conducted in accordance with the Declaration of Helsinki and approved by the IRB of the Dana-Farber/Harvard Cancer Center (DF/HCC Protocols: 11-035, 11-104, 17-000, 17-482; 09-204, 05-246). Patients with breast cancer who had CLIA-certified, tumor-only, exome-targeted next-generation sequencing (OncoPanel)^13^ successfully performed on primary or metastatic samples from July 2013 to December 2020 were included in this study. Patients with IBC who underwent OncoPanel testing were identified from the DFCI IBC Program database (Protocol 11-035) exported in December 2020, with final inclusion of those with both manually curated, sample-level clinicopathological information (n=140) and genomic results at time of metastatic disease^14^. IBC cases were diagnosed via American Joint Committee on Cancer (AJCC) guidelines as part of the DFCI IBC program. Cases without an IBC diagnosis were identified from the same database for comparison and designated non-IBC cases (n = 2317). American Society of Clinical Oncology/College of American Pathologists (ASCO/CAP) criteria were used to categorize each case into a breast cancer subtype—either HR-positive, HER2-positive, or TNBC—using molecular data at the time of metastatic diagnosis or, if no metastatic diagnostic biopsy was performed or data were unavailable, at the time of primary breast cancer diagnosis. In cases where patients were HR-positive and HER2-positive, they were classified as HER2-positive disease and all HR-positive classified cases were HER2-negative by ASCO/CAP criteria.

### Targeted tumor-only sequencing and tissue processing

Formalin-fixed, paraffin-embedded (FFPE) tumor tissue with >20% tumor cellularity per histopathologic review underwent DNA extraction and was assessed using OncoPanel (Supplemental Table S1)—a targeted, tumor-only sequencing platform that interrogates 277 (V1), 302 (V2), or 447 (V3) cancer-associated genes. The assay is performed centrally within the Clinical Laboratory Improvement Amendments (CLIA)-certified Center for Advanced Molecular Diagnostics at Brigham and Women’s Hospital (Boston, MA). Existent data was analyzed as part of the aforementioned consented research protocols.

### OncoPanel Analysis

Somatic alterations including SNVs CNVs were called as previously described^15, 16^. Given tumor-only data, common germline variants present in the gnomAD^17^ or Benign or Likely Benign variants in ClinVar^18^ databases were removed, unless also present in COSMIC^19^. To ensure consistent, unbiased calls across OncoPanel versions, genes that overlapped between the 3 different versions of OncoPanel were used for enrichment analyses of IBC vs. non-IBC (Supplemental Table S2). Somatic SNVs were further classified by their suspected oncogenicity using the OncoKB database and the predicted functional consequence of the mutation (i.e. nonsense, missense or frameshift mutations—Supplemental Table S3). For gene-level CNV calls, high amplifications and deep deletions were called as previously described (Supplemental Table S4)^13^. Sample-level TMB was estimated by dividing the frequency of all mutations in an individual by total panel size; units reflected as mutations/MB (Supplemental Table S5).

### Statistical Considerations

Clinicopathologic characteristics between IBC and non-IBC cases were compared using chi-squared for categorical and Wilcoxon test for continuous variables respectively. Regarding enrichment of genomic alterations in IBC vs non-IBC, such as SNVs and CNVs, cases were classified as either altered or unaltered based on the presence of a mutation or gene-level CNV. Enrichment for a particular group was performed using Fisher’s exact tests (Supplemental Tables S6-S9). If the variable of interest was continuous (i.e. TMB), a Wilcoxon test was used. For multiple comparisons, false-discovery rate (FDR) correction was performed using the Benjamini-Hochberg method to reduce the chance of Type I errors^20^. For the subtype-informed enrichment analysis, modeling was performed using a multivariate logistic regression accounting for HER2 and HR status. Only models that reached significance under multiple hypothesis correction for rejecting the log-likelihood null were included, as well as those that converged after 500 iterations. Only oncogenic mutations and high amplifications or deep deletions in over 1.5% of all IBC or non-IBC samples were included in this analysis (Supplemental Tables S10 – S11). For pathway alteration analysis, gene alterations were collated into 6 canonical cancer pathways (CELL_CYCLE, NOTCH, PI3K, RTK_RAS, TP53, WNT)—pathways were limited to those that contained at least 3 representative genes on the panel (Supplemental Table S12)^21^. The frequency of samples that harbored at least one alteration in each pathway was determined and compared between IBC and non-IBC cases (Supplemental Table S13). An exploratory survival analysis was performed using time from OncoPanel result date to last follow-up. Kaplan Meier analysis was performed in R (*survminer*) with statistical significance between survival curves assessed using the log-rank test.

### Data availability

The targeted panel sequencing data are continually deposited as part of the American Association for Cancer Research (AACR) Project GENIE—which is a publicly accessible cancer registry of clinicogenomic data from multiple institutions, of which DFCI is a contributing member. These data can be accessed after registering for Project GENIE and agreeing to the AACR’s terms of access (https://genie.cbioportal.org/login.jsp). Sample and gene-level mutation data can be found in Supplementary Data. Additional deidentified clinical information may be obtained upon request from the corresponding author and approval by the DF/HCC Breast Clinical Data and Biospecimens Users Committee—assuming adherence to and compatibility with the referenced protocols and local IRBs.

## RESULTS

### Clinicopathologic features of IBC vs. non-IBC

A total of 2457 patients were identified consisting of 140 cases of clinically confirmed IBC and 2317 cases of non-IBC (**Table 1**). Consistent with the more aggressive nature of IBC, patients with IBC were diagnosed with metastatic disease at an earlier age—developing metastases at a median age of 51 years versus 54 years of age in patients with non-IBC (p = 0.04). Patients with IBC notably had nearly double the rate of *de novo* metastases than patients with non-IBC disease (54.3% vs. 24.3%, p < 0.0001) with higher proportion of grade 3 histopathology at diagnosis (74.3% vs. 47.8%, p < 0.0001). As previously reported, IBC cases were enriched in hormone receptor (HR)-negative disease (54.3% vs. 25.1%, p < 0.0001) as well as HER2-positive disease (35.0% vs. 14.8%, p < 0.0001) yet interestingly harbored a lower proportion of HER2-low disease (IHC 1-2+ and fluorescence in situ hybridization (FISH)-negative; 17.9% vs 27.4%). Regarding the profiled samples—1622 were from metastatic disease (66.0% of total cohort, 51.4% of IBC cases, 67.0% of non-IBC cases), 770 were primary tumors (31.3% total, 48.6% IBC, 30.2% non-IBC), and 65 were representative of a local recurrence (2.6% total, all non-IBC cases).

**Table 1.** Patient and Clinical Characteristics Among Patients with Metastatic Breast Cancer with and without Inflammatory Breast Cancer (n=2457)

|  | Total Population<br>(n=2457) | Patients with IBC<br>(n=140) | Patients without IBC<br>(n=2317) | P-Value |
| --- | --- | --- | --- | --- |
| **Age in Years at time of metastatic diagnosis\*,**<br>**median (min, max)** | 54 (18, 91) | 51 (25, 91) | 54 (18, 89) | 0.04 |
| **Gender, n (%)** |  |  |  | 0.25 |
| Female | 2435 (99.1) | 140 (100) | 2295 (99.0) |  |
| Male | 22 (0.9) | 0 (0) | 22 (0.9) |  |
| **Race, n (%)** |  |  |  | 0.32 |
| African American | 116 (4.7) | 9 (6.4) | 107 (4.6) |  |
| American Indian, Aleutian, Eskimo | 2 (0.1) | 0 (0) | 2 (0.1) |  |
| Asian or Pacific Islander | 88 (3.6) | 4 (2.9) | 84 (3.6) |  |
| Caucasian | 2147 (87.4) | 126 (90.0) | 2021 (87.2) |  |
| Other | 49 (2.0) | 0 (0) | 49 (2.1) |  |
| Unknown | 55 (2.2) | 1 (0.7) | 54 (2.3) |  |
| **Ethnicity, n (%)** |  |  |  | 0.15 |
| Non-Spanish/Non-Hispanic | 2244 (91.3) | 134 (95.7) | 2110 (91.1) |  |
| Spanish/Hispanic | 87 (3.5) | 3 (2.1) | 84 (3.6) |  |
| Unknown | 126 (5.1) | 3 (2.1) | 123 (5.3) |  |
| **Stage at Initial Diagnosis, n (%)** |  |  |  | <0.0001 |
| DCIS | 28 (1.1) | 0 (0) | 28 (1.2) |  |
| I | 341 (13.9) | 0 (0) | 341 (14.7) |  |
| II | 812 (33.0) | 0 (0) | 812 (35.0) |  |
| III | 613 (24.9) | 64 (45.7) | 549 (23.7) |  |
| IV | 639 (26.0) | 76 (54.3) | 563 (24.3) |  |
| Unknown | 24 (1.0) | 0 (0) | 24 (1.0) |  |
| **Histology at Initial Diagnosis, n (%)** |  |  |  | 0.06 |
| DCIS | 34 (1.4) | 0 (0) | 34 (1.5) |  |
| Invasive Ductal | 1811 (73.7) | 107 (76.4) | 1704 (73.5) |  |
| Invasive Lobular | 311 (12.7) | 8 (5.7) | 303 (13.1) |  |
| Micropapillary | 3 (0.1) | 0 (0) | 3 (0.1) |  |
| Mixed (IDC & ILC) | 200 (8.1) | 18 (12.9) | 182 (7.8) |  |
| Mucinous | 9 (0.4) | 0 (0) | 9 (0.4) |  |
| Other | 16 (0.6) | 2 (1.4) | 14 (0.6) |  |
| Tubular | 3 (0.1) | 0 (0) | 3 (0.1) |  |
| Unknown | 70 (2.8) | 5 (3.6) | 65 (2.8) |  |
| **Histologic Grade at Initial Diagnosis, n (%)** |  |  |  | <0.0001 |
| Low | 142 (5.8) | 0 (0) | 142 (6.1) |  |
| Intermediate | 936 (38.1) | 32 (22.9) | 904 (39.0) |  |
| High | 1212 (49.3) | 104 (74.3) | 1108 (47.8) |  |
| Unknown | 167 (6.8) | 4 (2.9) | 163 (7.0) |  |
| **Hormone Receptor Status of Sample Tested,**<br>**n (%)** |  |  |  | <0.0001 |
| HR Positive | 1530 (62.3) | 59 (42.1) | 1471 (63.5) |  |
| HR Negative | 657 (26.7) | 76 (54.3) | 581 (25.1) |  |
| HR Not Done/Unknown | 270 (11.0) | 5 (3.6) | 265 (11.4) |  |
| **Estrogen Receptor Status of Sample Tested,**<br>**n (%)** |  |  |  | <0.0001 |
| ER Positive | 1399 (56.9) | 48 (34.3) | 1351 (58.3) |  |
| ER Low Positive | 93 (3.8) | 4 (2.8) | 89 (3.8) |  |
| ER Negative | 692 (28.2) | 83 (59.3) | 609 (26.3) |  |
| ER Not Done/Unknown | 273 (11.1) | 5 (3.6) | 268 (11.6) |  |
| **HER2 Status of Sample Tested, n(%)** |  |  |  | <0.0001 |
| HER2 Positive | 391 (15.9) | 49 (35.0) | 342 (14.8) |  |
| HER2 Negative | 1757 (71.5) | 87 (62.1) | 1670 (72.1) |  |
| HER2 Not Done/Unknown | 309 (12.6) | 4 (2.9) | 305 (13.1) |  |
| **HER2 Status (with Low Positive) of Sample**<br>**Tested, n(%)** |  |  |  | <0.0001 |
| HER2-Positive | 390 (15.9) | 48 (34.3) | 342 (14.8) |  |
| HER2-Low (IHC 1-2+, FISH-negative) | 661 (26.9) | 25 (17.9) | 636 (27.4) |  |
| HER2-0 (IHC 0) | 797 (32.4) | 45 (32.1) | 752 (32.5) |  |
| Not Done/Unknown | 609 (24.8) | 22 (15.7) | 587 (25.3) |  |
| **Type of Specimen Tested, n(%)** |  |  |  | <0.0001 |
| Primary Breast | 768 (31.3) | 68 (48.6) | 700 (30.2) |  |
| Local Recurrence | 65 (2.6) | 0 (0) | 65 (2.8) |  |
| Metastasis | 1624 (66.1) | 72 (51.4) | 1552 (67.0) |  |
| **Oncopanel Version, n(%)** |  |  |  | 0.5 |
| V1 | 194 (7.9) | 8 (5.7) | 186 (8.0) |  |
| V2 | 825 (33.6) | 45 (32.1) | 780 (33.7) |  |
| V 3 and 3.1 | 1429 (58.2) | 87 (62.1) | 1342 (57.9) |  |
| Unknown | 9 (0.4) |  | 9 (0.4) |  |

**IBC**, inflammatory breast cancer; **DCIS**, ductal carcinoma in situ; **IDC**, invasive ductal carcinoma; **ILC**, invasive lobular carcinoma; **HR**, hormone receptor; **ER**, estrogen receptor; **HER2**, human epidermal growth factor receptor 2; **FISH**, fluorescence in situ hybridization

### LumB-like histopathology enriched in HR-positive IBC

Given the aggressiveness of the disease, we determined if Luminal B tumors were more enriched in HR-positive/HER2-negative IBC vs non-IBC; inferred from histopathological data. Although a transcriptional subtype, we designated ‘LumB-inferred’ HR-positive tumors as those with grade 3 histology or progesterone receptor staining < 10% as these features have been associated with Luminal B tumors^22^. We observed a significant enrichment of LumB-inferred tumors in HR-positive IBC vs. non-IBC (70.0% vs. 40.1%, p = 0.0002).

### Landscape of somatic alterations in IBC

To expand beyond clinicopathologic correlates, the landscape of somatic alterations in IBC across subtypes was determined from targeted, tumor-only DNA panel sequencing. The most recurrent genomic alterations spanning all subtypes in IBC were those involving *TP53* (72%), *ERBB2* (32%), *PIK3CA* (24%), *CCND1* (12%), *MYC* (9%), *FGFR1* (8%) and *GATA3* (8%) (**Figure 1**). Among the most frequently altered genes, *TP53* single nucleotide variants (SNVs) showed the highest absolute frequency difference among all alterations between IBC and non-IBC cases (**Figure 2A**)—particularly in HER2-positive and HR-positive disease—with an alteration frequency of 85.1% in IBC vs. 62.7% in non-IBC and 50.0% vs. 26.8%; respectively. *ESR1* and *GATA3* alteration frequencies were, as expected, present predominantly in HR-positive disease—with an alteration frequency of 2.5% in IBC vs. 12.0% in non-IBC for *ESR1* and 15.0% vs. 12.9% for *GATA3*. Regarding copy number variants (CNVs), the most recurrent alterations were similar to those previously reported and included amplifications of regions involving *MYC*, *ERBB2*, *FGFR1* as well as deletions in *CDKN2A/B* (**Figure 2B**).

**Figure 1.**
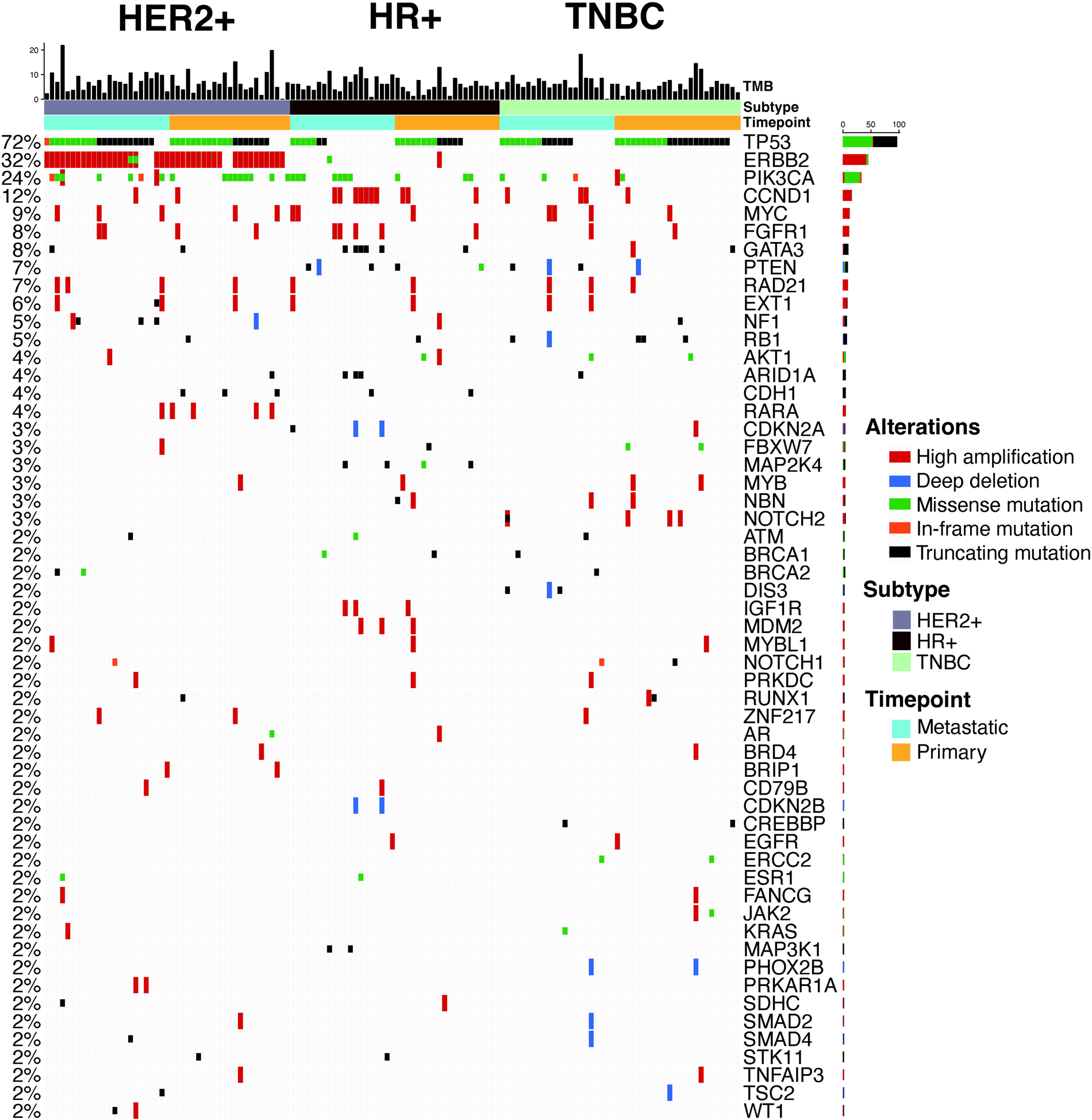
Genomic landscape of inflammatory breast cancers. Genes ordered by percentage of somatic alterations in overall cohort. Samples divided by breast cancer subtype and subdivided by primary or metastatic tissue tested. All variants represent oncogenic mutations or deep deletions/high amplifications. Tumor mutational burden (TMB) (mut/mb) is recorded on the top barplot of the OncoPrint. **HER2+,** human epidermal growth factor receptor 2-positive; **HR+,** hormone receptor-positive; **TNBC,** triple-negative breast cancer

**Figure 2.**
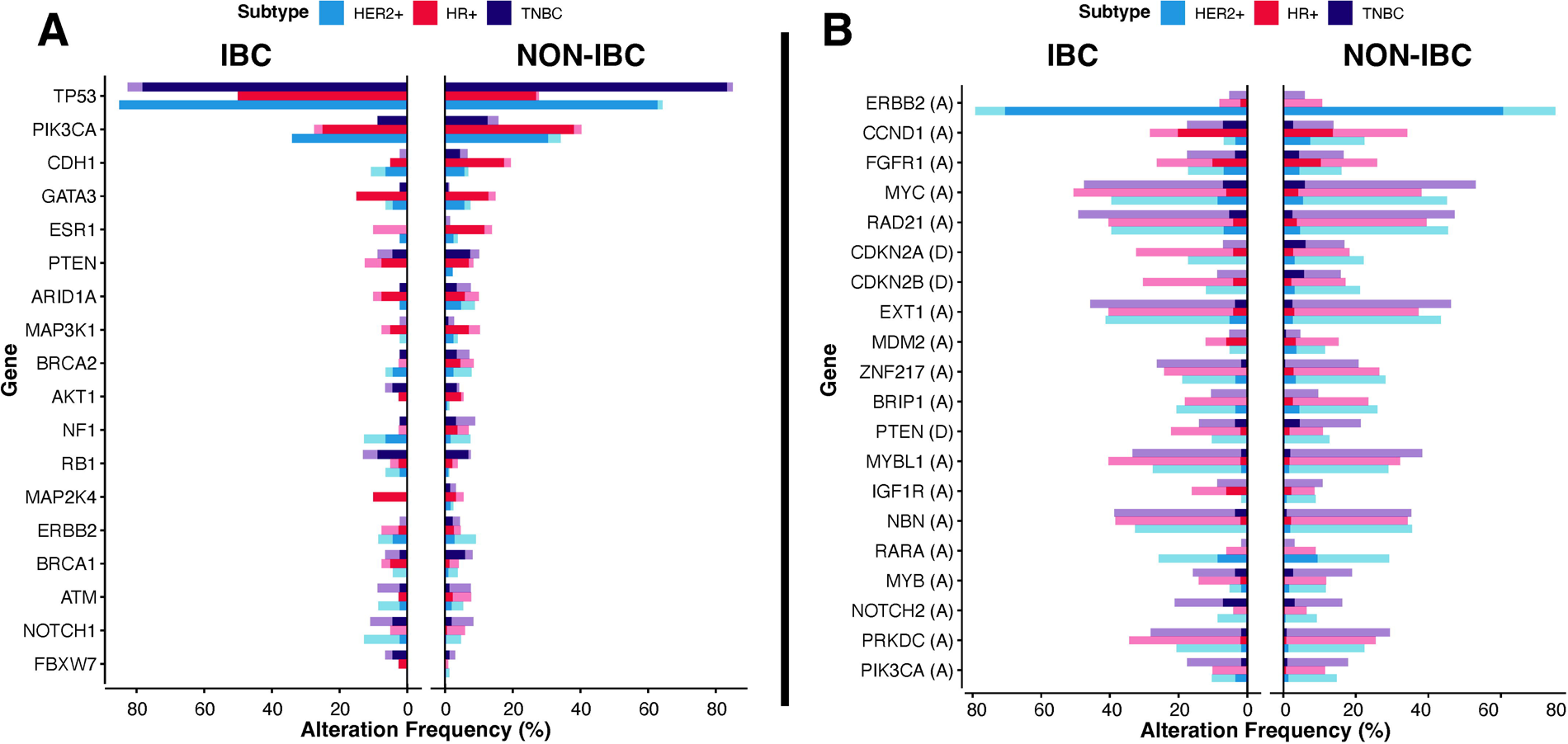
Frequency of most common SNVs (A, left) and CNVs (B, right) in IBC and non-IBC colored by subtype. Shading represents the percentage of oncogenic events (defined by OncoKB for SNVs, defined by estimated high amplification or predicted double copy deletion for CNVs). For Figure 2B, an annotation of “(A)” beside a gene represents an amplification and “(D)” represents a deletion. **IBC,** inflammatory breast cancer; **<u>SNVs,</u>** single nucleotide variants; **CNVs,** copy number variants

### Limited somatic differences between IBC vs non-IBC, except for a significant enrichment in TP53 alterations

To determine statistically significant, subtype-informed enrichments in IBC vs. non-IBC, we implemented a logistic regression analysis to account for HR and HER2 status. Among the most recurrent SNV alterations, only *TP53* mutations were significantly enriched after accounting for multiple hypothesis testing (odds ratio (OR), 2.10 [95% confidence interval (CI), 1.36 – 3.25, adjusted p-value = 0.016) (**Figure 3A**). No significant, gene-level enrichments were observed in CNVs in IBC (**Figure 3B**). When assessing alterations in an analysis of six canonical cancer pathways (CELL_CYCLE, NOTCH, PIK3, RTK_RAS, TP53, WNT – **Figure 4**)— the most significantly altered pathways enriched in IBC by subtype were members of the TP53 pathway in both HER2-positive (Frequency of alterations: 89.3% vs. 74.9%, p-value = 0.03) and HR-positive disease (60.0% vs. 41.6%, p-value = 0.02) disease and members of the NOTCH signaling pathway in HER2-positive disease (27.6% vs 14.7%, p-value = 0.03). No nominally significant enrichments were observed in triple-negative breast cancer (TNBC).

**Figure 3.**
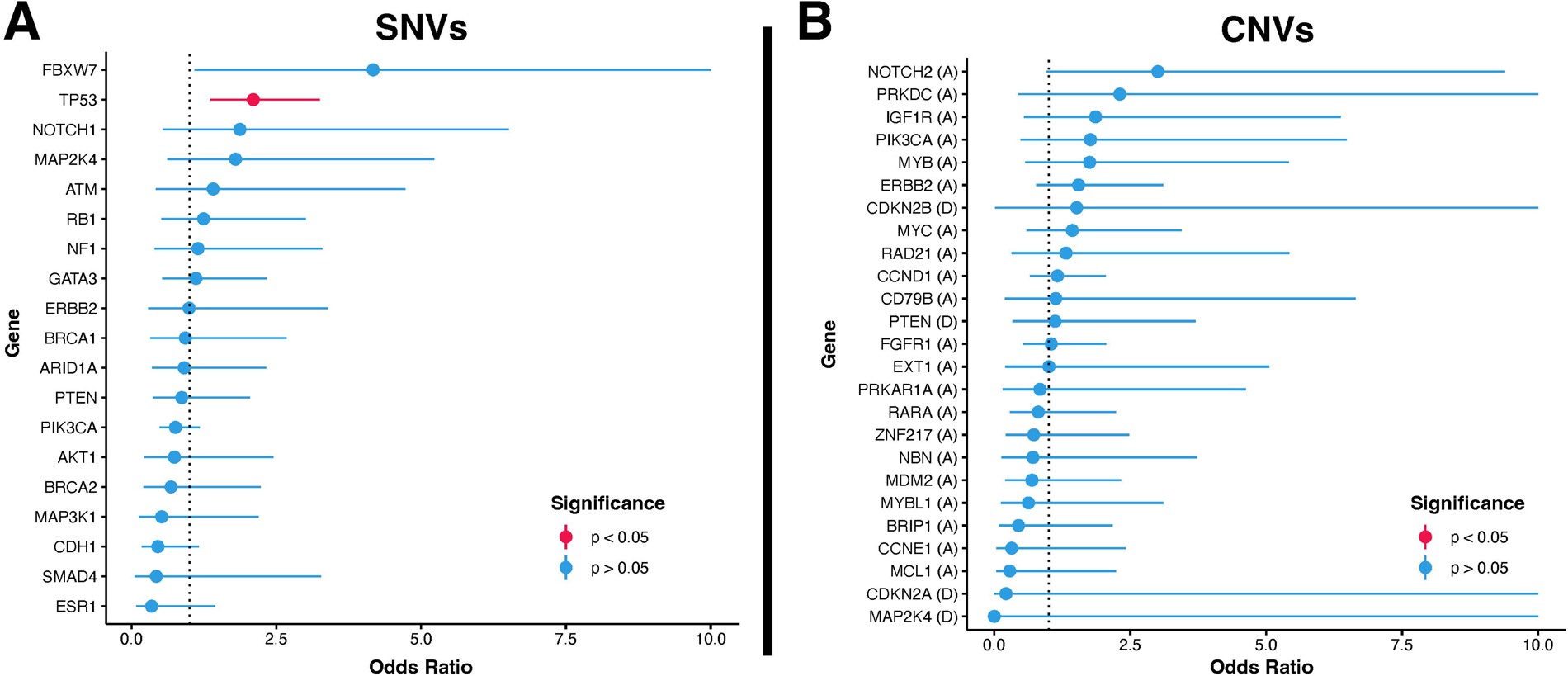
Enrichment analysis of SNVs (A, left) and CNVs (B, right) in IBC. Modeling performed using multivariate logistic regression accounting for HER2 and HR status. Only models that converged after 500 iterations are shown. Oncogenic mutations and high amplifications or deep deletions that appeared in over 1.5% of either all IBC or non-IBC samples were included in the analysis. **IBC,** inflammatory breast cancer; **SNVs,** single nucleotide variants; **CNVs,** copy number variants

**Figure 4.**
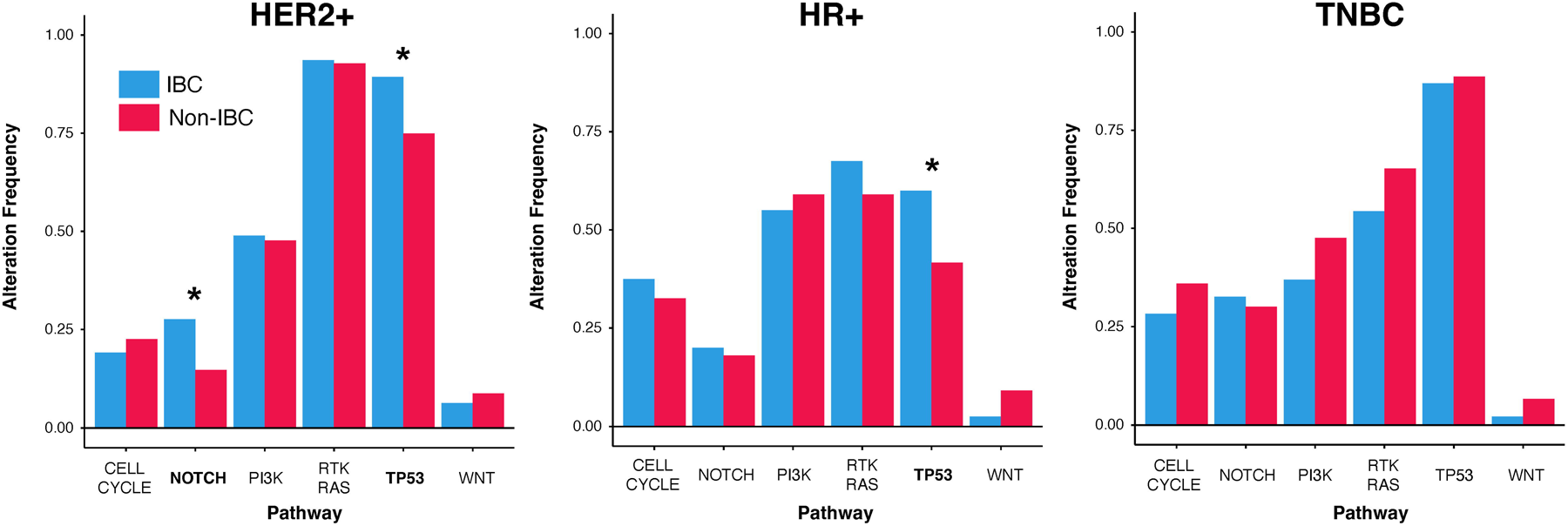
Comparison of somatic alterations grouped by biological pathways between IBC and non-IBC cases. Proportion of samples with alterations within 6 biological pathways, segregated by breast cancer subtype; colored by IBC status (blue = IBC, red = non-IBC). Nominally significant enrichment (p < 0.05) highlighted with * above bar plots. **HER2+,** human epidermal growth factor receptor 2-positive; **HR+,** hormone receptor-positive; **TNBC,** triple-negative breast cancer

### Lack of tumor mutational burden differences in IBC vs non-IBC

Tumor mutational burden (TMB) was then assessed in IBC and non-IBC cases. Again, minimal differences were observed when comparing IBC and non-IBC—segregated by subtype or by primary vs. metastatic disease (**Figure 5**). The IBC vs. non-IBC median TMB were as follows by subtype: **HR-positive disease** 6.05 [Q1 3.8, Q3 7.79] vs 6.08 [Q1 4.56, Q3 8.47], **HER2-positive disease** 6.84 [Q1 3.8, Q3 9.68] vs 6.08 [Q1 3.98, Q3 8.47], **TNBC** 6.05 [Q1 4.56, Q3 7.26] vs 6.64 [Q1 4.56, Q3 8.47].

**Figure 5.**
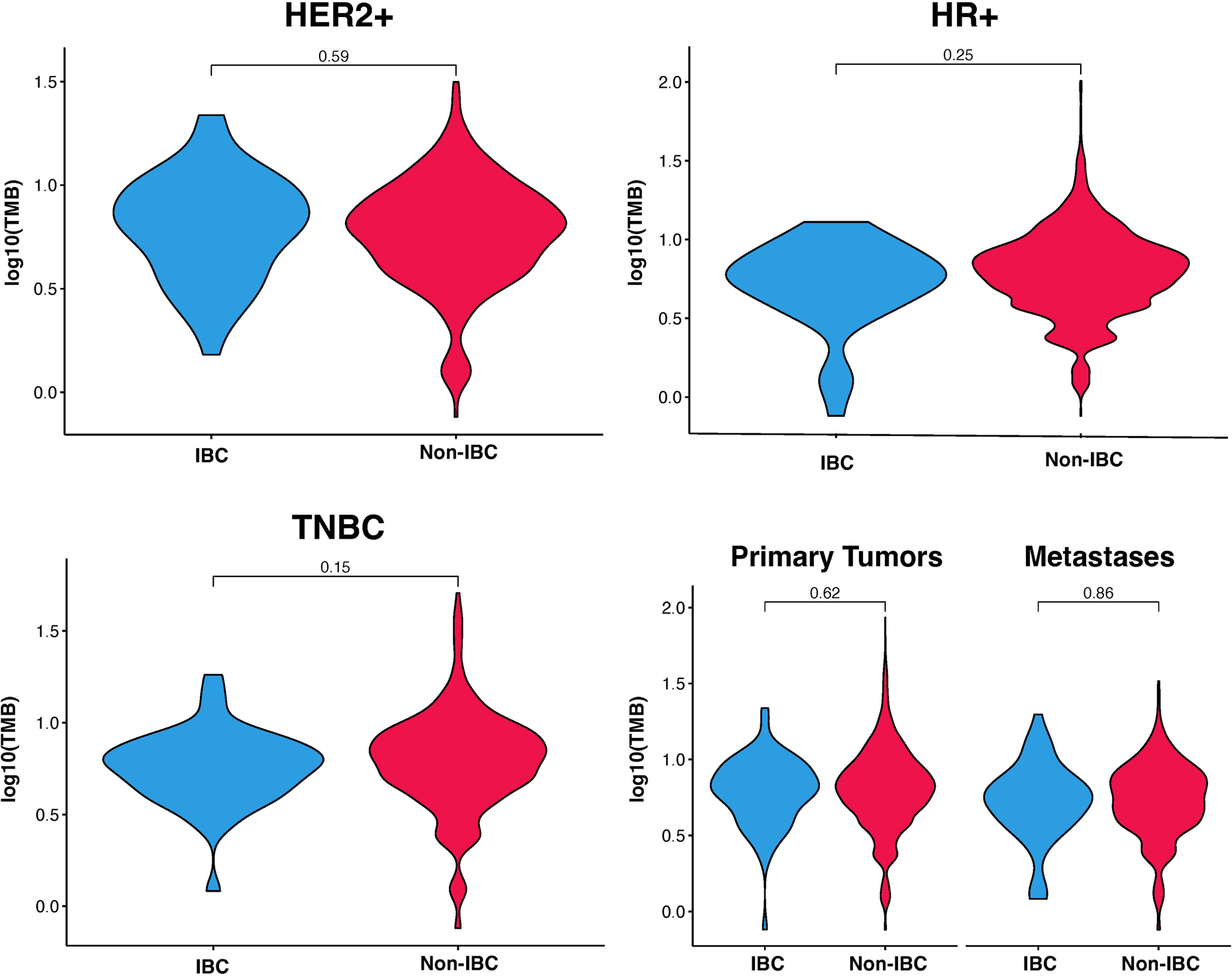
Comparison of TMB between IBC and non-IBC cases. Tumor mutational burden (TMB, mutations / MB) between IBC and non-IBC cases; divided by subtype. Bottom right plot shows tumors segregated by primary vs. metastatic lesion assayed. **IBC,** inflammatory breast cancer; **HER2+,** human epidermal growth factor receptor 2-positive; **HR+,** hormone receptor-positive; **TNBC,** triple-negative breast cancer

#### Landscape of TP53 alterations in IBC and association with worse outcomes in HR-positive disease

Albeit at a higher frequency, no identifiable difference in mutation patterns was observed in IBC and non-IBC across p53; with the majority being predicted loss-of-function alterations (Figure 6A). An exploratory analysis of survival outcomes showed *TP53* alterations were associated with worse outcomes in HR+ IBC (logrank p-value = 0.028) with a median overall survival—following metastatic genomic tumor testing—of 495 days in *TP53* mutated cases versus 993 days in cases without a TP53 mutation detected (Figure 6B). No significant survival differences were observed in HER2+ disease when segregated by *TP53* mutation status (logrank p-value = 0.48).

**Figure 6.**
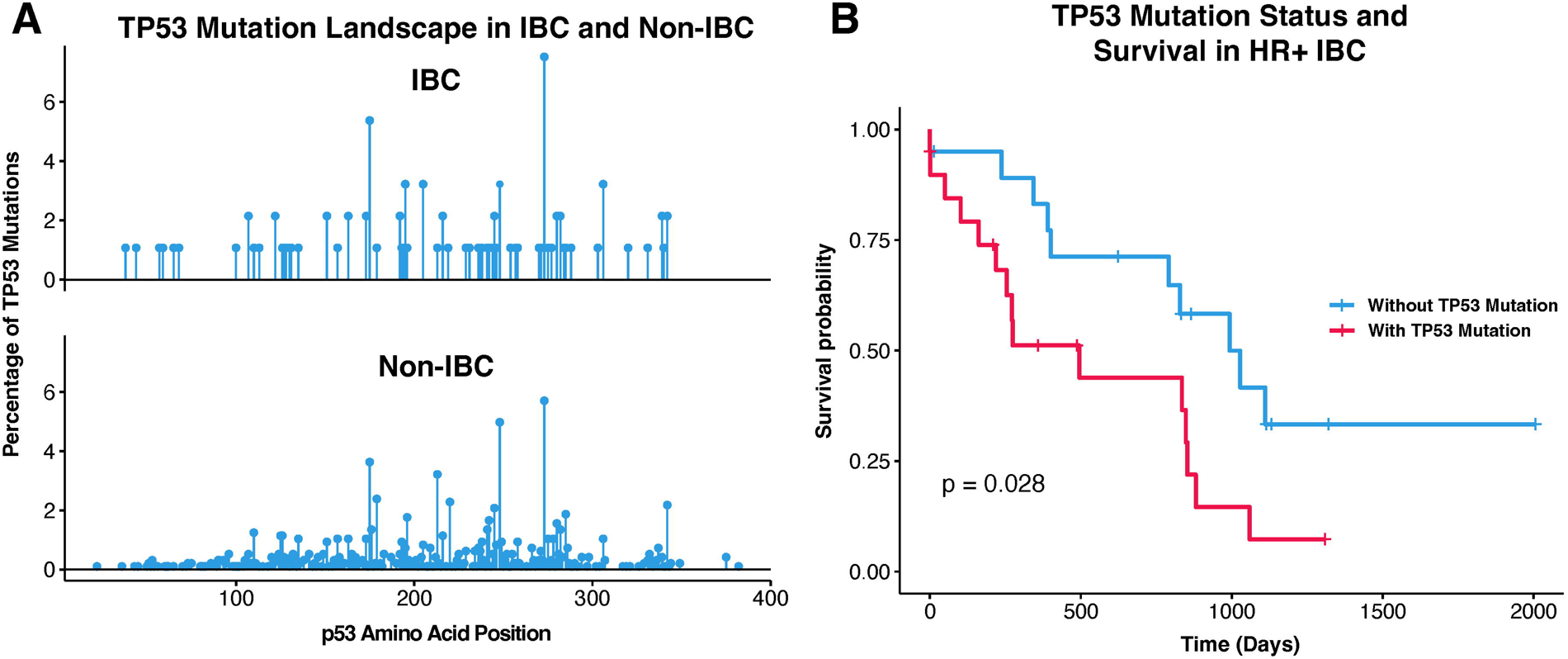
Landscape of TP53 alterations in IBC and association with worse outcomes in HR+ IBC. **(A)** Lollipop plot of TP53 mutations identified in IBC cases (top) and non-IBC cases (bottom). **(B)** Overall survival after OncoPanel testing in advanced IBC cases segregated by presence or absence of TP53 mutation. Median overall survival (date of OncoPanel testing to date of last follow-up) 495 days in *TP53* mutated cases versus 993 days in cases without a TP53 mutation detected. Logrank p-value shown on plot.

## DISCUSSION

IBC stands out as a unique and notably aggressive variant of breast cancer, marked by distinct clinical features and unfavorable outcomes. Despite its distinctiveness, a thorough understanding of this disease remains elusive—largely due to its rarity and the challenge of assembling large cohorts. In this study, we perform one of the most comprehensive clinicogenomic analyses of IBC to date, all conducted within a dedicated IBC center, which enabled a comprehensive and subtype-informed analysis of clinicopathological characteristics and associated genomic data that are enriched in IBC vs non-IBC.

In summary, our clinicopathological findings are consistent with prior reports— with an enrichment of higher risk features such as increased frequency of *de novo* metastases, higher grade tumors with an inferred LumB-like histopathology, and a younger age at metastatic diagnosis. Genomically, we find a significant enrichment in *TP53* alterations—particularly in HR-positive and HER2-positive disease—with a frequency of 50.0% and 85.1% respectively. This study did not identify other somatic alterations unique to IBC when correcting for molecular subtypes. We also observed a lack of gene-level CNV enrichments in IBC and a similar overall TMB when compared to non-IBC cases. This data suggests IBC has limited unique genomic features versus non-IBC, at least based on this limited targeted panel sequencing.

The most frequent somatic alterations in IBC involve *TP53*, *ERBB2*, *PIK3CA*, *CCND1*, *MYC*, *FGFR1*, *GATA3* and *PTEN*—which is generally quite consistent with prior studies on IBC and overall matches the distribution of driver alterations in unselected cohorts of breast cancer, like the TCGA^23–25^. When accounting for subtype, no significant enrichments were observed in IBC either in our gene-level CNV or SNV analysis except for a significant enrichment in somatic *TP53* alterations. Increased frequency of mutations in *TP53* have been found in smaller studies, yet we notably did not observe statistically significant enrichments in other genes previously reported to be enriched in IBC, including *ERBB2* mutations, when performing our subtype-informed analyses^26^. We also found no differences in TMB, which contrasts some prior studies that were performed exclusively on primary tumors^24, 26^. This could be somewhat explained by our inclusion of tumors representative of more advanced disease^25^, although even when segregating by primary and metastases, we did not find a statistically significant difference in TMB between IBC and non-IBC cases. Notably, NOTCH signaling has been implicated in IBC pathogenesis and we did observe a nominally significant increased frequency of alterations in NOTCH pathway—specifically in HER2-positive disease^27^.

*TP53* mutations were found to be significantly enriched in IBC—especially in subtypes not thought to harbor these mutations as frequently, such as in the HR-positive setting where we observe a *TP53* alteration frequency of 50% in IBC vs. 27.8% in non-IBC. Additionally, these alterations were associated with worse outcomes in HR+ IBC. As has been widely studied, *TP53* is broadly classified as a tumor suppressor encoding for a transcription factor with frequent loss-of-function somatic mutations in human cancers; which can confer increased cellular proliferation via cell cycle dysregulation, inhibit DNA damage repair processes, and lead to decreased apoptosis among many other reported functions^28–31^. Recently, several small molecules that restore p53 function through various mechanisms are advancing through clinical development, challenging the historical notion that *TP53* may be undruggable^32^. Given the frequency of TP53 mutations we observe in IBC, a question would be whether some of these small molecules or strategies targeting the *TP53* axis might be particularly suitable for preclinical testing in models of IBC—especially in the HR+ setting given the poorer overall survival we observe in our exploratory analysis. Biologically, reports have shown that loss of p53 in breast cancer drives metastasis through WNT-mediated recruitment of pro-metastatic systemic inflammation and neutrophilia in mouse models^33^—which may be one possible hypothesis connecting the frequency of *TP53* mutations to some of the high-risk features seen clinically in IBC; such as a higher frequency of *de novo* metastatic disease. *TP53* has also been implicated specifically in progression and metastasis through other mechanisms, such as facilitating epithelial to mesenchymal transition, cell motility, as well as pro-metastatic receptor tyrosine kinase signaling^34^. We postulate that further attention should be made preclinically to determine which of these mechanisms, if any, may be playing a role in the progression of IBC and whether modifying the *TP53* axis therapeutically could serve as a novel approach for IBC-directed therapy.

Besides p53 and potential NOTCH enrichments in HER2-positive disease, limited somatic differences between IBC and non-IBC were observed. This conclusion is somewhat limited by our use of targeted panel testing and perhaps more comprehensive assessments of the genome may yield more differences. However, a recent study that employed whole-genome sequencing to profile IBC cases (n=20) also did not reveal many somatic enrichments in IBC vs non-IBC—including non-coding alterations—other than *MAST2*; albeit this cohort was relatively small^11^. Collectively, our data suggest limited distinct alterations in the IBC genome even with a larger cohort and when a subtype-informed analysis is performed—at least in commonly interrogated, cancer-related genes.

A major limitation of this study includes the targeted nature of the sequencing panel. Perhaps a more comprehensive analysis utilizing whole exome or genome sequencing may reveal coding and non-coding alterations enriched in IBC. Also, although the ability to detect SNVs and CNVs with targeted-sequencing data is somewhat robust, more nuanced structural variation (complex rearrangements, chromothripsis, chromoplexy, etc.) and RNA-level changes (fusions, splice variants, expression-based enrichments, etc.) cannot be interrogated by the sequencing platform in this study. Lastly, our analysis included many patients that harbored metastatic disease at the time of targeted tumor sequencing testing, which may introduce some selection bias versus other studies.

Given a lack of clear IBC specific biomarkers at the somatic level when correcting for subtype, other features should be studied to explain what may be driving the unique IBC clinical phenotype. This work further supports the notion that the genomic landscape of IBC may not be distinct from that of non-IBC except for *TP53* mutations and perhaps NOTCH signaling alterations. Moving forward, understanding the pathogenesis of IBC may demand discovery efforts using features not captured by standard genomic profiling—such as environmental exposures, germline-somatic influences, RNA-level alterations, and microenvironmental interactions—as well as the application of novel genomic technologies such as single-cell sequencing and spatial profiling^35^.

## ACKNOWLEDGMENTS

The authors would like to thank the patients who contributed samples and their time to help make this study possible.

## PREVIOUS PRESENTATION

Nolan Priedigkeit, Beth Harrison, Melissa Hughes, Robert Shue, Yvonne Li, Greg Kirkner, Marie Claire Remolano, Sarah Strauss, Janet Files, Anne-Marie Feeney, Ayesha Mohammed-Abreu, Ana Garrido-Castro, Romualdo Barroso Sousa, Brittany Bychkovsky, Faina Nakhlis, Jennifer R. Bellon, Tari King, Bruce E. Johnson, Lynette Sholl, Deborah Dillon, Beth Overmoyer, Sara M. Tolaney, Andrew Cherniack, Nancy Lin, Filipa Lynce. (December 5-9, 2023) Comprehensive clinicogenomic characterization of inflammatory breast cancer. Presented at the 2023 San Antonio Breast Cancer Symposium, San Antonio, TX, USA.

## AUTHOR CONTRIBUTIONS

Study concept and design (NP, BH, RS, MH, YL, BO, SMT, ADC, NUL, FL) acquisition, analysis, or interpretation of data (all authors); drafting of the manuscript (all authors); critical revision of the manuscript for important intellectual content (all authors); administrative, technical, or material support (all authors).

## References

1 Robertson FM, Bondy M, Yang W et al. Inflammatory breast cancer: the disease, the biology, the treatment. CA Cancer J Clin 2010; 60 (6): 351–375.

2 Lim B, Woodward WA, Wang X et al. Inflammatory breast cancer biology: the tumour microenvironment is key. Nat Rev Cancer 2018; 18 (8): 485–499.

3 Hance KW, Anderson WF, Devesa SS et al. Trends in inflammatory breast carcinoma incidence and survival: the surveillance, epidemiology, and end results program at the National Cancer Institute. Journal of the National Cancer Institute 2005; 97 (13): 966–975.

4 Kertmen N, Babacan T, Keskin O et al. Molecular subtypes in patients with inflammatory breast cancer; a single center experience. J BUON 2015; 20 (1): 35–39.

5 Dano D, Lardy-Cleaud A, Monneur A et al. Metastatic inflammatory breast cancer: survival outcomes and prognostic factors in the national, multicentric, and real-life French cohort (ESME). ESMO Open 2021; 6 (4): 100220.

6 Garrido-Castro AC, Regan MM, Niman SM et al. Clinical outcomes of de novo metastatic HER2-positive inflammatory breast cancer. NPJ Breast Cancer 2023; 9 (1): 50.

7 Bonnier P, Charpin C, Lejeune C et al. Inflammatory carcinomas of the breast: a clinical, pathological, or a clinical and pathological definition? Int J Cancer 1995; 62 (4): 382–385.

8 Anderson WF, Schairer C, Chen BE et al. Epidemiology of inflammatory breast cancer (IBC). Breast Dis 2005; 22: 9–23.

9 Van Laere SJ, Ueno NT, Finetti P et al. Uncovering the molecular secrets of inflammatory breast cancer biology: an integrated analysis of three distinct affymetrix gene expression datasets. Clin Cancer Res 2013; 19 (17): 4685–4696.

10 Ross JS, Ali SM, Wang K et al. Comprehensive genomic profiling of inflammatory breast cancer cases reveals a high frequency of clinically relevant genomic alterations. Breast Cancer Res Treat 2015; 154 (1): 155–162.

11 Li X, Kumar S, Harmanci A et al. Whole-genome sequencing of phenotypically distinct inflammatory breast cancers reveals similar genomic alterations to non-inflammatory breast cancers. Genome Med 2021; 13 (1): 70.

12 Hester RH, Hortobagyi GN, Lim B. Inflammatory breast cancer: early recognition and diagnosis is critical. Am J Obstet Gynecol 2021; 225 (4): 392–396.

13 Garcia EP, Minkovsky A, Jia Y et al. Validation of OncoPanel: A Targeted Next-Generation Sequencing Assay for the Detection of Somatic Variants in Cancer. Arch Pathol Lab Med 2017; 141 (6): 751–758.

14 Hughes M, Frank E, Merrill M et al. Abstract P4-10-04: EMBRACE (Ending metastatic breast cancer for everyone): A comprehensive approach to improve the care of patients with metastatic breast cancer. Cancer Research 2018; 78 (4_Supplement): P4-10-04-P14-10-04.

15 Garrido-Castro AC, Spurr LF, Hughes ME et al. Genomic Characterization of de novo Metastatic Breast Cancer. Clin Cancer Res 2021; 27 (4): 1105–1118.

16 Tarantino P, Gupta H, Hughes ME et al. Comprehensive genomic characterization of HER2-low and HER2-0 breast cancer. Nat Commun 2023; 14 (1): 7496.

17 Karczewski KJ, Francioli LC, Tiao G et al. The mutational constraint spectrum quantified from variation in 141,456 humans. Nature 2020; 581 (7809): 434–443.

18 Landrum MJ, Lee JM, Benson M et al. ClinVar: improving access to variant interpretations and supporting evidence. Nucleic Acids Res 2018; 46 (D1): D1062–D1067.

19 Tate JG, Bamford S, Jubb HC et al. COSMIC: the Catalogue Of Somatic Mutations In Cancer. Nucleic Acids Res 2019; 47 (D1): D941–D947.

20 Benjamini Y, Hochberg Y. Controlling the False Discovery Rate: A Practical and Powerful Approach to Multiple Testing. Journal of the Royal Statistical Society: Series B (Methodological) 1995; 57 (1): 289–300.

21 Sanchez-Vega F, Mina M, Armenia J et al. Oncogenic Signaling Pathways in The Cancer Genome Atlas. Cell 2018; 173 (2): 321–337 e310.

22 Cancello G, Maisonneuve P, Rotmensz N et al. Progesterone receptor loss identifies Luminal B breast cancer subgroups at higher risk of relapse. Ann Oncol 2013; 24 (3): 661–668.

23 Koboldt DC, Fulton RS, McLellan MD et al. Comprehensive molecular portraits of human breast tumours. Nature 2012; 490 (7418): 61–70.

24 Liang X, Vacher S, Boulai A et al. Targeted next-generation sequencing identifies clinically relevant somatic mutations in a large cohort of inflammatory breast cancer. Breast Cancer Res 2018; 20 (1): 88.

25 Richard F, De Schepper M, Maetens M et al. Comparison of the genomic alterations present in tumor samples from patients with metastatic inflammatory versus non-inflammatory breast cancer reveals AURKA as a potential treatment target. Breast 2023; 69: 476–480.

26 Matsuda N, Lim B, Wang Y et al. Identification of frequent somatic mutations in inflammatory breast cancer. Breast Cancer Res Treat 2017; 163 (2): 263–272.

27 Bertucci F, Rypens C, Finetti P et al. NOTCH and DNA repair pathways are more frequently targeted by genomic alterations in inflammatory than in non-inflammatory breast cancers. Mol Oncol 2020; 14 (3): 504–519.

28 Olivier M, Hollstein M, Hainaut P. TP53 mutations in human cancers: origins, consequences, and clinical use. Cold Spring Harb Perspect Biol 2010; 2 (1): a001008.

29 Giono LE, Manfredi JJ. The p53 tumor suppressor participates in multiple cell cycle checkpoints. J Cell Physiol 2006; 209 (1): 13–20.

30 Muller PA, Vousden KH. p53 mutations in cancer. Nat Cell Biol 2013; 15 (1): 2–8.

31 Levine AJ. The many faces of p53: something for everyone. J Mol Cell Biol 2019; 11 (7): 524–530.

32 Hassin O, Oren M. Drugging p53 in cancer: one protein, many targets. Nat Rev Drug Discov 2023; 22 (2): 127–144.

33 Wellenstein MD, Coffelt SB, Duits DEM et al. Loss of p53 triggers WNT-dependent systemic inflammation to drive breast cancer metastasis. Nature 2019; 572 (7770): 538–542.

34 Tang Q, Su Z, Gu W et al. Mutant p53 on the Path to Metastasis. Trends Cancer 2020; 6 (1): 62–73.

35 Moffitt JR, Lundberg E, Heyn H. The emerging landscape of spatial profiling technologies. Nat Rev Genet 2022; 23 (12): 741–759.

